# Robust CENP-A incorporation in human cells is independent of transcription and cohesin components

**DOI:** 10.1101/2025.04.29.651290

**Authors:** Reito Watanabe, Carlos Perea-Resa, Michael D. Blower

## Abstract

Centromeres are essential chromosomal components that ensure proper cell division by serving as assembly sites for kinetochores, which connect chromosomes to spindle microtubules. Centromeres are marked by the evolutionarily conserved centromere-specific histone H3 variant, CENP-A, which is deposited into centromere nucleosomes during G1 in human cells. Centromeres retain cohesin, a ring-like protein complex during mitosis, protecting sister chromatid cohesion and centromere transcription to prevent chromosome missegregation. Previous work in *Drosophila* has suggested that centromere transcription and centromeric RNAs are important for CENP-A deposition in chromatin. During mitosis centromeric cohesin is critical for centromere transcription. However, it is not clear how or if centromeric transcription and cohesin contribute to CENP-A deposition in G1 in human cells. To address these questions, we combined a cell synchronization strategy with the Auxin Inducible Degron technology and transcription inhibition in human cells. In contrast to *Drosophila cells*, our results demonstrated that neither centromeric transcription nor cohesin is required for CENP-A deposition in human cells. Our data demonstrate clear differences in the CENP-A deposition mechanism between human and *Drosophila* cells. These findings provide deeper insights into the plasticity underlying centromere maintenance and highlight evolutionary divergence in centromere maintenance systems across species.

## Introduction

Chromosomes are equally distributed to daughter cells during mitosis through attachment of sister chromatids to the spindle microtubules by the kinetochore, which is assembled on centromeric DNA (Fukagawa & Earnshaw, 2014; McKinley & Cheeseman, 2016). Centromeres are epigenetically defined by the presence of the evolutionarily conserved centromere-specific histone H3 variant, Centromere protein A, CENP-A, which is deposited into centromere nucleosomes during G1 in human cells (Black et al., 2004, 2007; Blower & Karpen, 2001; Earnshaw & Rothfield, 1985; Foltz et al., 2006; Fujita et al., 2007; Henikoff et al., 2000; Jansen et al., 2007; Meluh et al., 1998; Palmer et al., 1991; Régnier et al., 2003; Sekulic et al., 2010). Artificial tethering of CENP-A to noncentromeric chromatin induced de novo centromeres in many organisms, which were epigenetically maintained after the tethering was removed (Barnhart et al., 2011; Bassett et al., 2012; C.-C. Chen et al., 2014; Dunleavy et al., 2009; Foltz et al., 2009; Heun et al., 2006; Hori et al., 2012). In addition, CENP-A is required for faithful chromosome segregation and for the proper localization of most kinetochore proteins to centromeres (Foltz et al., 2006; Hayashi et al., 2004; Howman et al., 2000; Oegema et al., 2001). Thus, a significant question is how CENP-A deposits into centromere nucleosomes. Previously, work showed that the Mis18 complex, composed of Mis18α, Mis18β, and M18BP1/KNL2 (KNL2), recruits HJURP-CENP-A to centromeres in many vertebrates (Dambacher et al., 2012; Dunleavy et al., 2009; Erhardt et al., 2008; Foltz et al., 2009; Hayashi et al., 2004; Hori et al., 2017; Jansen et al., 2007; Maddox et al., 2007; Moree et al., 2011; Shuaib et al., 2010). The process is tightly regulated by phosphorylation during the cell cycle to ensure that CENP-A loading occurs in G1 (Flores Servin et al., 2023; French & Straight, 2019a; McKinley & Cheeseman, 2014; Müller et al., 2014; Pan et al., 2017; Silva et al., 2012; Stankovic et al., 2017; Wang et al., 2014; Yu et al., 2015). DNA replication dilutes chromatin-bound CENP-A and these genomic sites are filled by placeholder conventional H3-containing nucleosomes. It is not yet clear how placeholder nucleosomes are removed from chromatin during CENP-A deposition in G1.

Transcription through centromeric chromatin has been proposed as a mechanism that could evict placeholder H3 nucleosomes from centromere chromatin. Recent work has demonstrated that centromeres are transcribed by RNA polymerase II (RNAP II) during the cell cycle in many different organisms (Blower, 2016; Catania et al., 2015; Chan et al., 2012; Chan & Wong, 2012; Choi et al., 2011; Ferri et al., 2009; Li et al., 2008; Liu et al., 2015; McNulty et al., 2017; Ohkuni & Kitagawa, 2011; Perea-Resa et al., 2020; Rošić et al., 2014; Wong et al., 2007). Centromere transcription is regulated by CENP-B, CENP-I, and CENP-C, which are constitutively associated with centromere chromatin (Bury et al., 2020; Y. Chen et al., 2021a; Hirai et al., 2022; Sikder et al., 2025). Centromeric RNAs are required for CENP-A maintenance, kinetochore assembly, and accurate chromosome segregation (Blower, 2016; Bobkov et al., 2018; Chan et al., 2012; Liu et al., 2015; McNulty et al., 2017; Perea-Resa et al., 2020; Rošić et al., 2014). Removal of H3K4me2 in Human Artificial Chromosomes (HAC) reduces local transcription, localization of HJURP, and CENP-A (Bergmann et al., 2011). In addition, the Facilitates Chromatin Transcription (FACT) complex, is essential for maintaining CENP-A in chickens and flies (C.-C. Chen et al., 2015; Deyter & Biggins, 2014; Okada et al., 2009). Collectively, these findings suggest that transcription may contribute to CENP-A maintenance or deposition. However, considering that centromere RNA mediates CENP-A incorporation and kinetochore formation, a short-term experimental system with specific transcription inhibition is needed to investigate the importance of transcription or chromatin remodeling itself in CENP-A incorporation. Work in *Drosophila* cells using transcription inhibitors has shown that transcription is vital for incorporating CID/dCENP-A into complete nucleosomes (Bobkov et al., 2018). On the other hand, a recent study suggested that transcription was not required for CENP-A incorporation during *Drosophila* embryogenesis (Ghosh & Lehner, 2022). From these conflicting observations, it is not clear whether transcription facilitates CENP-A incorporation at native centromeres.

During mitosis, centromere transcription is mediated by the retention of cohesin, a ring-like protein complex that protects physical cohesion between sister chromatids (Perea-Resa et al., 2020; Peters et al., 2008). Interestingly, a recent paper showed centromere transcription also mediated mitotic cohesion, suggesting a positive feedback mechanism between cohesin and transcription (Y. Chen et al., 2021b). Cohesin is a multiprotein complex comprising core subunits Smc1, Smc3, and Rad21, and either STAG1 or STAG2 (Arruda et al., 2020; Casa et al., 2020; Horsfield, 2023; Losada et al., 2000; Peters et al., 2008). Cohesin is important for extruding DNA into loops and may promote transcription through regulating enhancer promoter contacts. Because cohesin promotes mitotic centromere transcription it is possible that cohesin is involved in CENP-A loading in G1. In this work, focusing on human cells, we examine the functional importance of cohesin and transcription in CENP-A deposition with specific transcription inhibitors and a rapid protein degradation system.

## Results and Discussion

### Rapid depletion of cohesin components does not affect newly synthesized CENP-A localization to centromeres in human cells

To examine the importance of cohesin components for newly synthesized CENP-A localization in human cells, we combined the pulse chase CENP-A labeling assay with auxin-inducible degron 2 (AID2) system targeting cohesin components and M18BP1/KNL2 (KNL2) (Jansen et al., 2007; Nishimura et al., 2009; Yesbolatova et al., 2020), which is an essential factor of CENP-A recruitment (Supplemental Figure 1, A, and B). In brief, cells were synchronized at the G1/S boundary by thymidine block for 24 hours, and SNAP-Cell Block quenched nucleosomal CENP-A. Then, cells were released into nocodazole, and newly synthesized cytoplasmic CENP-A-SNAP was labeled with SNAP-TMR. We then removed SNAP-TMR and triggered protein degradation using addition of 5-Ph-IAA followed by measurement of centromeric CENP-A-SNAP-TMR in G1 cells. Rapid KNL2 depletion during mitosis to the G1 phase caused defects in the CENP-A localization consistent with previous reports (Figure 1, A and B, Supplemental Figure 1D).

**Figure 1.**
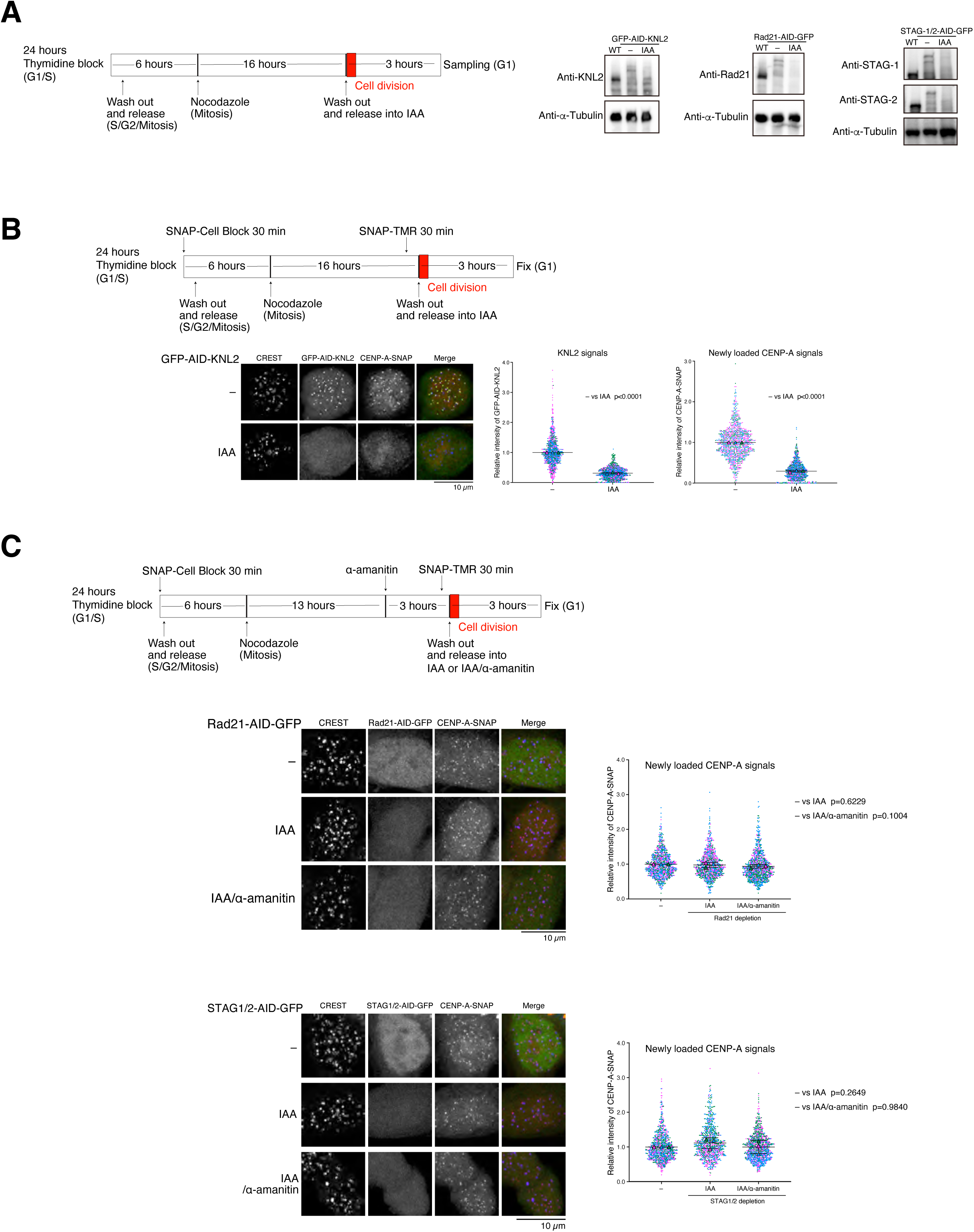
Cohesin components are not required for newly synthesized CENP-A localization in cells. (A) Schematic representation of the experiment (left). The protein levels of endogenous KNL2, Rad21, STAG1, and STAG2, which were tagged with AID and GFP in human DLD1 cells constitutively expressing TIR1-F74G (right). AID-tagged protein was conditionally depleted by 5-Ph-IAA (IAA) addition. (B) Schematic representation of the experiment (top). Localization of GFP-fused KNL2 (green) and newly deposited CENP-A-SNAP labeled with SNAP-TMR (red) during G1 phase 3 hours after the IAA addition. Centromeres were stained by CREST as a marker (blue). Images are max projected from z-stacks. GFP and TMR signals on centromeres were quantified. Each dots indicate a signal from one centromere. The Bar graph indicates the mean with SD from an average of three replicates. (C) Schematic representation of the experiment (top). Localization of GFP-fused Rad21 or STAG1/2 (green) and newly deposited CENP-A-SNAP was stained with SNAP-TMR (red) during G1 phase at 3 hours after the IAA addition or a combination of 6 hours of α-amanitin with IAA for 3 hours (IAA/α-amanitin). Centromeres were stained by CREST as a marker (blue). Images are max projected from z-stacks. TMR signals on centromeres were quantified. Each dots indicate a signal from one centromere. The Bar graph indicates the mean with SD from an average of three replicates.

Although Rad21 is a cohesin core component critical for DNA looping (Jeppsson et al., 2022), newly synthesized CENP-A was correctly localized onto centromeres in Rad21-depleted cells (Figure 1, A and C, Supplemental Figure 1, A, B, and D). This indicates that cohesin DNA-binding and looping activity were not required for newly synthesized CENP-A localization. The cohesin components, STAG1 or STAG2, stabilize transcription and R-loops independently of cohesin function (Porter et al., 2023). Thus, STAG proteins could mediate chromatin remodeling for CENP-A loading. Depletion of STAG1 or STAG2 alone did not show growth defects, consistent with previous papers (Lelij et al., 2020; Mondal et al., 2019)(Supplemental Figure 1, C and D). Therefore, we established STAG1 and STAG2 AID-tagged cells for rapid double depletion, which resulted in a severe growth defect (Figure 1 A, Supplemental Figure 1, A, B, and D). Double depletion of STAG 1 and 2 did not alter CENP-A localization (Figure 1 C), demonstrating that cohesin and cohesin-mediated chromatin remodeling were not required for newly synthesized CENP-A localization in human cells.

We then examined the function of transcription-dependent chromatin remodeling in the CENP-A loading context in human cells. Recent work in *Drosophila* cells and the early embryo showed that inhibition of transcription did not impair centromere localization of CID/dCENP-A and CAL1, the CENP-A loading machinery (Bobkov et al., 2018; Ghosh & Lehner, 2022). We established an AID-based CENP-A loading system in human cells expressing AID-GFP-tagged CENP-A from one endogenous allele. Cells were incubated in IAA for 2 days to remove chromatin-bound CENP-A-GFP prior to the thymidine block. Following thymidine washout CENP-A-AID-GFP was newly synthesized and localized in the G1, but not during mitosis (Figure 2 A, Supplemental Figure 2, A and B). We established this system as an alternative to the CENP-A-SNAP loading system because it has a much higher signal-to-noise ratio in G1 cells. A recent paper indicated that α-amanitin, but not triptolide, inhibits centromere transcription in human cells (Y. Chen et al., 2021a). Thus, cells were treated with triptolide or α-amanitin, which both decreased RNA pol II CTD phospho-Ser2. However, they did not alter CENP-A localization (Figure 2 A, Supplemental Figure 3). We also analyzed KNL2 and CENP-A in cells expressing KNL2-AID-GFP and CENP-A-SNAP, which were treated with transcription inhibitors. Transcription inhibition did not affect the localization of CENP-A-SNAP or KNL2 to centromeres in G1, consistent with data from *Drosophila* (Figure 2 B). We conclude that chromatin remodeling mediated by transcription is not required for CENP-A and KNL2 localization in human cells.

**Figure 2.**
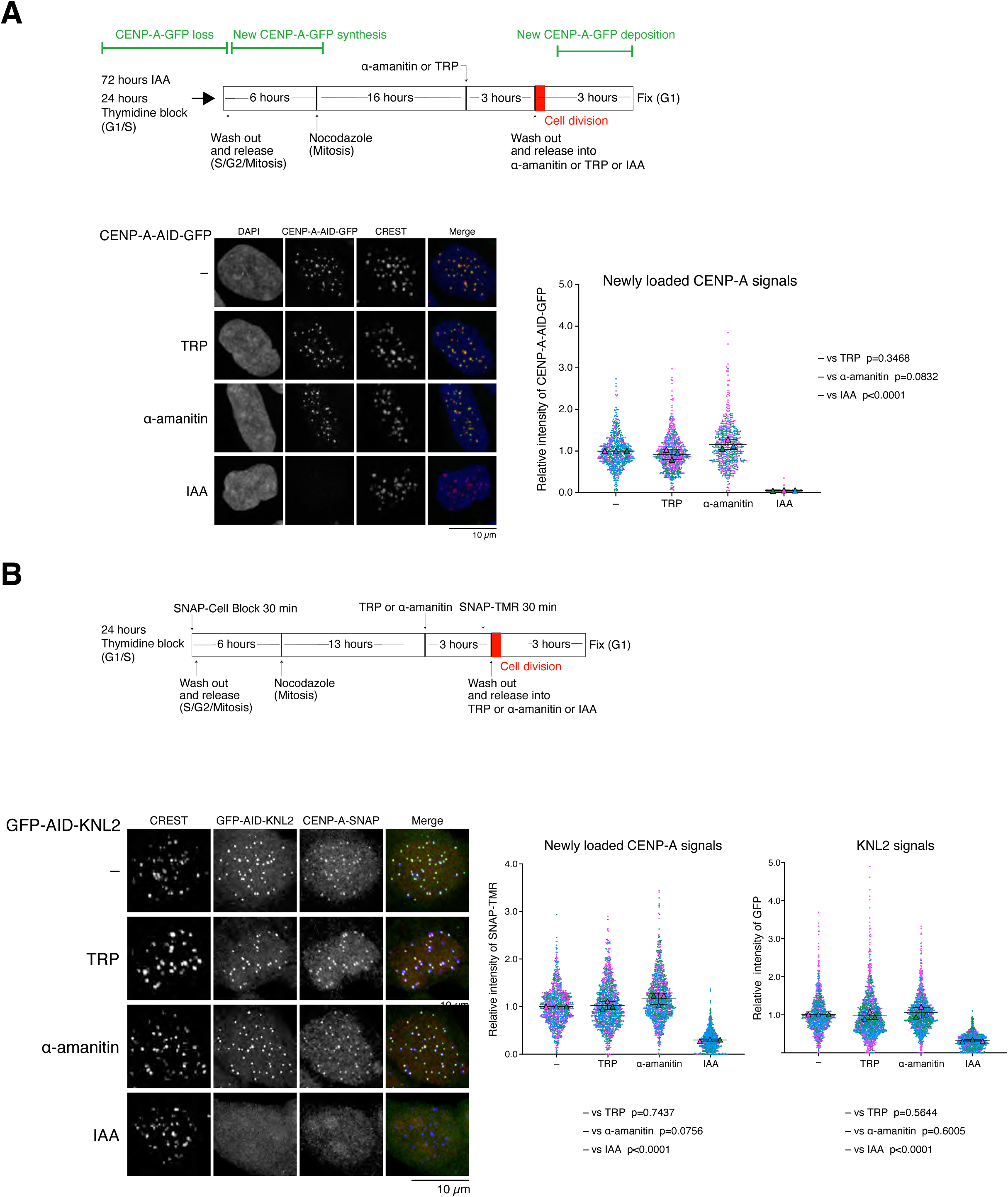
KNL2 and newly synthesized CENP-A localization do not depend on transcription in human cells. (A) Schematic representation of the experiment (top). Localization of newly deposited CENP-A-AID-GFP (green) during G1 phase at 6 hours after Triptolide (TRP) or α-amanitin, or 3 hours after the IAA addition in HeLa cells. DNA was stained with DAPI (blue). Centromeres were stained by CREST as a marker (red). Images are max projected from z-stacks. GFP signals on centromeres were quantified. Each dot indicates a signal from one centromere. The Bar graph indicates the mean with SD from an average of three replicates. (B) Schematic representation of the experiment (top). Localization of newly deposited CENP-A-SNAP (red) and GFP-AID-KNL2 (green) during G1 phase at 6 hours after Triptolide (TRP) or α-amanitin, or 3 hours after the IAA addition in DLD1 cells. Centromeres were stained by CREST as a marker (blue). Images are max projected from z-stacks. GFP and TMR signals on centromeres were quantified. Each dots indicate a signal from one centromere. The Bar graph indicates the mean with SD from an average of three replicates.

Transcription inhibition blocked CENP-A incorporation into centromere nucleosomes in *Drosophila* cells, but not CENP-A localization to centromeres, implying that CENP-A incorporation into nucleosomes depends on transcription-coupled chromatin remodeling (Bobkov et al., 2018). We hypothesized that chromatin remodeling, depending on DNA looping by cohesin, might be required for CENP-A incorporation in conditions where transcription is silenced. However, depletion of cohesin components, combined with α-amanitin, did not affect CENP-A localization, suggesting chromatin remodeling by both cohesin and transcription was not required for CENP-A localization (Figure 1 C).

### Centromeric localization of CENP-A and KNL2 is stable following a combination of transcriptional inhibition and pre-extraction

A previous *Drosophila* study showed that transcription stabilized CENP-A in nucleosomes by removing unincorporated CENP-A on nucleosomes with high salt treatment before fixing cells (Bobkov et al., 2018). To evaluate whether the CENP-A loading mechanism in flies is conserved in human cells, newly synthesized CENP-A localization was examined in human cells treated with 0.5M NaCl or 0.5% Triton-X in the absence or presence of triptolide or α-amanitin (Figure 3 A, Supplemental Figure 4). Under any conditions, CENP-A localization was not impaired, suggesting that CENP-A was incorporated into nucleosomes even when transcription is silenced in human cells. To further investigate the stability of the upstream factor of CENP-A incorporation under transcription inhibition, both CENP-A and KNL2-GFP signals were analyzed with 0.5M NaCl under transcription inhibition. Salt treatment combined with transcription inhibition did not evict CENP-A or KNL2 localization in human cells (Figure 3 B), indicating that KNL2 is tightly bound to centromere chromatin. This contrasts with *Drosophila* CAL1, which was excluded from chromatin by salt treatment (Bobkov et al., 2018).

**Figure 3.**
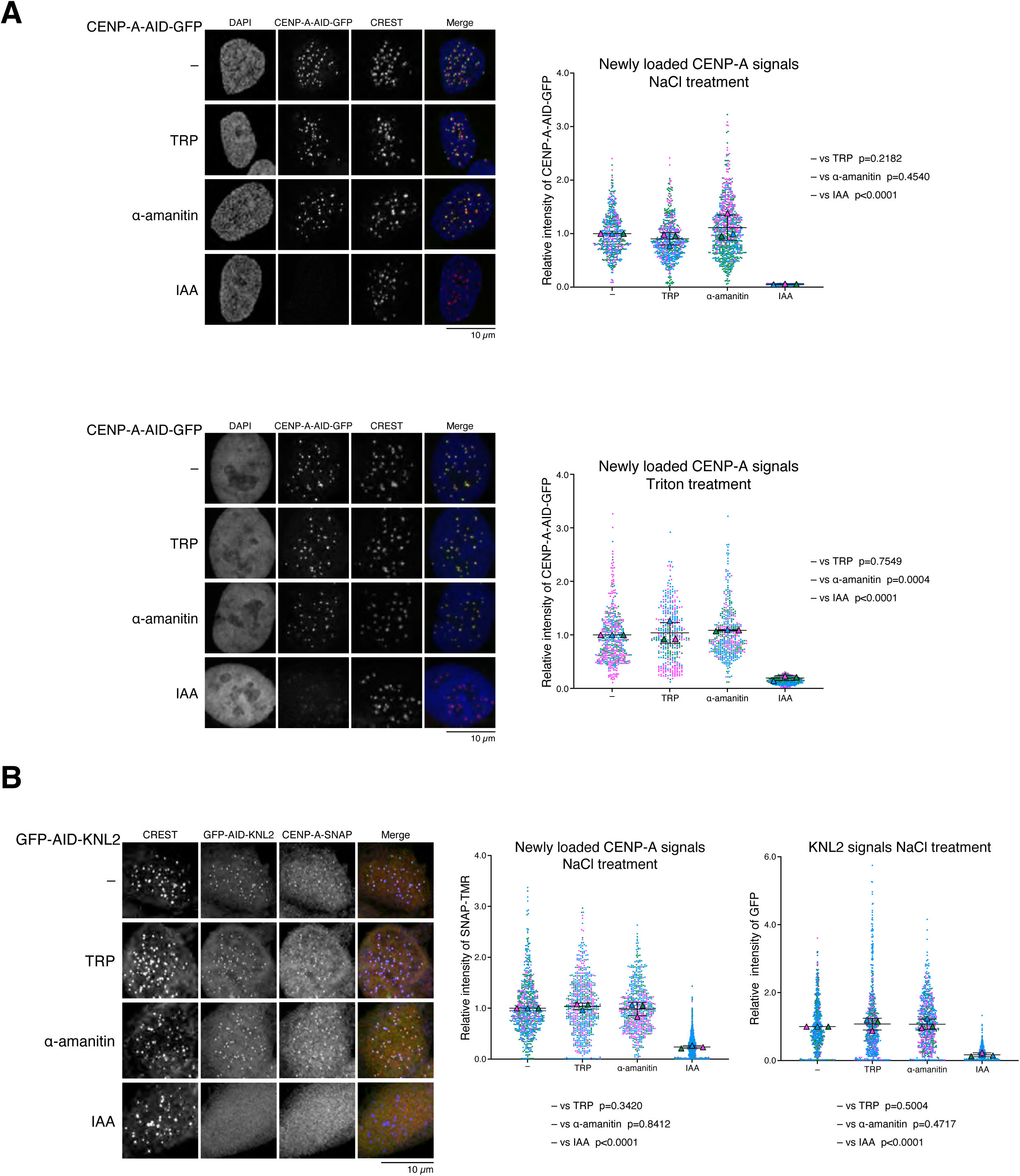
CENP-A deposition and KNL2 localization are stable in salt extraction and detergent treatment in pre-fixed cells incubated with transcription inhibitors. (A) Localization of newly deposited CENP-A-AID-GFP (green) during G1 phase at 6 hours after Triptolide (TRP) or α -amanitin, or 3 hours after the IAA addition in HeLa cells, as shown in the schematic representation of the experiment in Fig. 2A. Cells were fixed in formaldehyde after 30 min of 0.5 M NaCl-PBS salt extraction (top) or 60 s 0.5% Triton-X-PBS treatment (bottom). DNA was stained with DAPI (blue). Centromeres were stained by CREST as a marker (red). Images are max projected from z-stacks. GFP signals on centromeres were quantified. Each dots indicate a signal from one centromere. The Bar graph indicates the mean with SD from an average of three replicates. (B) Localization of newly deposited CENP-A- SNAP (red) and GFP-AID-KNL2 (green) during G1 phase at 6 hours after Triptolide (TRP) or α-amanitin, or 3 hours after the IAA addition in DLD1 cells as shown in the schematic representation of the experiment in Fig. 2B. Cells were fixed in formaldehyde after 30 min of 0.5 M NaCl-PBS salt extraction. Centromeres were stained by CREST as a marker (blue). Images are max projected from z-stacks. GFP and TMR signals on centromeres were quantified. Each dots indicate a signal from one centromere. The Bar graph indicates the mean with SD from an average of three replicates.

### CENP-A incorporation into the nucleosome did not depend on transcription

Our results indicated that KLN2 and newly-synthesized CENP-A were stably bound to nucleosomes without transcription. However, this did not address whether KNL2 could tether CENP-A to chromatin in the absence of transcription-coupled chromatin incorporation (Figure 4A). To test if KNL2 is tethering CENP-A to chromatin in the absence of transcription we examined the effects of KNL2 depletion on CENP-A centromere localization following transcription inhibition. (Figure 4 A). Mitotic cells are released into the G1 phase with α-amanitin for 2 hours, then cells are incubated with IAA for 1 hour to degrade KNL2 rapidly. If KNL2 is tethering CENP-A to chromatin in the absence of transcription we predicted that CENP-A localization would be lost following KNL2 depletion. CENP-A was localized to centromeres after KNL2 degradation in control cells, demonstrating that release from nocodazole for 2 hours was sufficient for CENP-A recruitment onto centromeres (Figure 4 A and B). CENP-A also localized normally in KNL2 depleted cells in which transcription is inhibited. Collectively these data suggest that transcription is not required for stable CENP-A incorporation into chromatin in G1.

**Figure 4.**
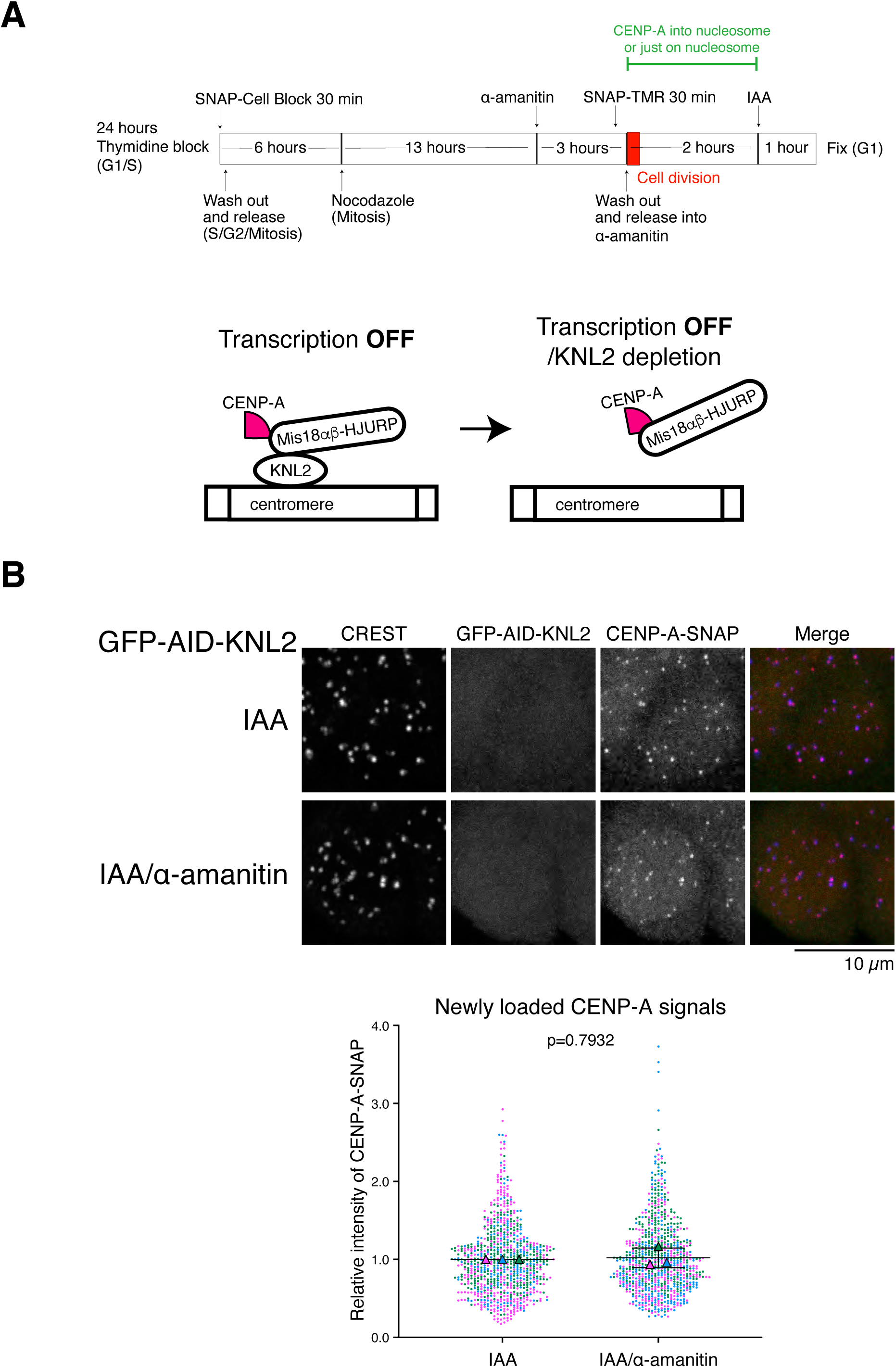
CENP-A was deposited into nucleosomes even in the presence of transcription inhibitors and became independent of KNL2. (A) Schematic representation of the experiment (top). Schematic representation of the model for transcription inhibitor blocks CENP-A deposition into nucleosomes that depends on KNL2. (B) Localization of newly deposited CENP-A-SNAP (red) and GFP-AID-KNL2 (green) during G1 phase presence or absence of α-amanitin for 6 hours, and 3 hours after the IAA addition. Centromeres were stained by CREST as a marker (blue). Images are max projected from z-stacks. TMR signals on centromeres were quantified. Each dots indicate a signal from one centromere. The Bar graph indicates the mean with SD from an average of three replicates.

Here, we demonstrated that human CENP-A localization and chromatin incorporation did not require cohesin or transcription. Our observations suggest that the CENP-A deposition mechanism in human cells is distinct from that of *Drosophila*, which relies on transcription. In *Drosophila*, CID/dCENP-A directly binds to CAL1, which interacts with the constitutive centromere protein, CENP-C (C.-C. Chen et al., 2014; Phansalkar et al., 2012; Schittenhelm et al., 2010; Unhavaithaya & Orr-Weaver, 2013). In contrast, human CENP-A interacts with CENP-C through M18BP1/KNL2 and HJURP in the Mis18α/β complex (Cao et al., 2024; Pan et al., 2019; Shuaib et al., 2010; Tachiwana et al., 2015; Thamkachy et al., 2024). The varied requirement for transcription in CENP-A incorporation between different species might reflect differences in protein structure of the CENP-A loading machinery, the cell cycle stage of CENP-A loading, or differences in the protein composition of the centromere/kinetochore complex (Barnhart et al., 2011; Barnhart-Dailey et al., 2017; Hu et al., 2023; McKinley & Cheeseman, 2014; Parashara et al., 2024).

It is noteworthy that transcription or its dependent chromatin remodeling is required for CENP-A maintenance in many organisms and several biological contexts. The functional disruption of transcription-associated chaperone FACT showed defective CENP-A assembly in chickens and flies (C.-C. Chen et al., 2015; Okada et al., 2009). Active transcription transiently expels centromere histones and Spt6 serves to replace CENP-A evicted by transcription (Bobkov et al., 2020). Centromeric transcription destabilized centromere nucleosomes in the prophase I-arrested starfish oocyte, to allow gradual replacement of new CENP-A, but was not required for canonical CENP-A recruitment in G1 (Swartz et al., 2019). Taken together with our data and published works, canonical CENP-A deposition in G1 does not depend on transcription in many organisms. However, centromeric transcription allows the replacement of CENP-A, which is crucial for maintaining proper CENP-A amount in quiescent cells, or in some specific organisms. Our findings highlight evolutionary divergence in centromere maintenance systems in different cell types and across species.

## MATERIALS AND METHODS

### Cell culture

DLD-1 cells were cultured in RPMI-1640 (Sigma) + 10% FBS (Cytivia) + Pen/Strep. HeLa cells were cultured in DMEM (Sigma) + 10% FBS (Cytivia) + Pen/Strep. To induce AID-tagged protein degradation, cells were treated with 1 µM 5-Ph-IAA for the indicated hours in figures. DLD-1 cells and HeLa cells were treated with 2 mM thymidine in complete medium for 24 hours, washed twice in complete medium, and released in complete medium for the indicated hours in the figures, followed by the addition of nocodazole to a final concentration of 100 ng/ml, and shacked off mitotic cells and washed out by complete medium three times and cultured on collagen covered cover grass for indicated hours in the figures. SNAP-Cell Block (NEB) and SNAP-TMR (NEB) were used at 2 µM for 30 min. Triptlide was used at 1 µM for the indicated times. α-amanitin was used at 10µg/ml for the indicated times. For drug selection, the transfected DLD-1 and HeLa cells were selected in medium containing 10 µg/ml of Blasticidin S HCL, 100 µg/ml of Zeosin, 500 µg/ml of G418, 1 µg/ml of puromycin, or 200 µg/ml of Hygromycin B.

### Cell lines

As shown in Supplemental Figure 1A, to constitutively express TIR1-F74G in DLD-1 or HeLa cells at the AAVS1 locus, pMB1398 (EF1-a promoter-TIR1-F74G-P2A- Blasticidin resistant gene) and pMB1422 (Cas9-D10A targeting AAVS locus) were transfected by Lipofectamine 3000 (Thermo). Each plasmid DNA (2.5 μg) and Lipofectamine 3000 were incubated in 500 µl of Opti-MEM at room temperature for 20 minutes and then added to the cells incubated in –Pen/Strep medium. Cells were selected with the proper drug for 2 weeks, 72 hours after transfection. Then, pMB1405 (CENP-A C-terminus HR with SNAP, PGK promoter-Zeosin resistant gene) and pMB1329 (Cas9 targeting CENP-A C-terminus) were transfected into DLD-1 cells expressing TIR1-F74G. After selection, we got DLD-1 cells expressing CENP-A-SNAP from a single allele, to express Rad21-mAID-GFP under control of the endogenous promoter, pMB 1446 (Rad21 C-terminus HR with mAID-GFP-P2A-Neomycin resistant gene), and pMB1153 (Cas9 targeting Rad21 C-terminus), were transfected. As with Rad21, to express STAG1 or STAG2 tagged with mAID-GFP, pMB1461 (STAG1 C-terminus HR with mAID-GFP-P2A-Neomycin resistant gene) and pMB1353 (Cas9 targeting STAG1 C-terminus), or pMB1440 (STAG2 C-terminus HR with mAID-GFP-P2A-Neomycin resistant gene) and pMB1354 (Cas9 targeting STAG2 C-terminus) were transfected. To express GFP-mAID-KNL2 under the control of the endogenous promoter, pMB1481 (KNL2 N-terminus HR with Hygromycin resistant gene-P2A-GFP-mAID) and pMB1486 (Cas9 targeting KNL2 N-terminus) were transfected. And to degrade both STAG1 and STAG2, pMB1506 (STAG2 C-terminus HR with mAID-GFP-P2A-Puromycin resistant gene) and pMB1354 (Cas9 targeting STAG2 C-terminus) were transfected into DLD-1 cells expressing STAG1-mAID-GFP. To express CENP-A-mAID-GFP in HeLa cells, pMB1408 (CENP-A C-terminus HR with mAID-GFP, PGK promoter-Zeosin resistant gene) and pMB1329 (Cas9 targeting CENP-A C-terminus) were transfected, and single allele knocking-in cells were selected. Knocking-in in all cell lines was confirmed by genome PCR and immunoblotting.

### Plasmid constructions

All plasmid constructs were created using Infusion cloning. We designed 500- 750bp homology regions flanking the target stop codon of Rad21, STAG1, STA2, CENP-A, and fused these to the mAID-GFP selection cassette using Infusion cloning. For the KNL2 N-terminus knocking-in plasmid, we thank Dr. Aaron Straight for providing plasmid including the homology regions flanking the target start codon of KNL2 (French & Straight, 2019b). We inserted the Hygromycin resistance cassette and P2A into the plasmid by infusion cloning. All plasmid sequences were confirmed by whole plasmid sequencing.

### Genotyping PCR

DLD-1 cells or HeLa cells were trypsinized and harvested, washed with PBS. Genome was provided and amplified by Platinum™ Direct PCR Universal Master Mix (Thermo A44647100). PCR primers are followings: For the AAVS locus, forward primer GCTAGTCTTCTTCCTCCAACCC, reverse primer CAAGCTCTCCCTCCCAGGATC. For the KNL2 locus, forward primer CACCGAAGAATTTACTTACCTCCAG, reverse primer AAACCTGGAGGTAAGTAAATTCTTC. For the Rad21 locus, forward primer CAAGCTATTGAGCTGACACAGG, reverse primer CCAGTGTTACTGATGGAAAGAAGTG. For the STAG1 locus, forward primer GAAGGGACATAATTCAGCCCGTAAC, reverse primer ATGCTCCTCTTCTGTCACTGC. For the STAG2 locus, CAAGGGAGAGTTGCTCCATTCATC, reverse primer TATTCAGGCAAACAAAACTGCC. For the CENP-A locus, forward primer ACCACCTATTTTCCACTGAACTTACC, reverse primer GTAACCAGAACATCAAAGCTTACAGG.

### Immunoblotting

DLD-1 cells were trypsinized and harvested, washed with PBS, and suspended in 1× LSB 1× Laemmli sample buffer (LSB; final 2 × 10^4 cells/µl) followed by sonication and heating for 5 min at 96°C. Proteins were separated on a 7.5% SDS-PAGE gel, handmade and transferred to Immobilon (Millipore) by Trans-Blot Turbo (BIORAD) with 2.5 A for 12 min. Membranes were washed with PBS + 2vol/vol% Tween 20 and blocked with PBS + 2vol/vol% Tween 20 supplemented with 5% skim milk for 20 min at room temperature. Then, membranes were washed with PBS + 2vol/vol% Tween 20 for 5 min three times and incubated with primary antibody in Western Blocker and Signal Enhancer (MP) overnight at 4°C. Then, membranes were washed with PBS + 2vol/vol% Tween 20 for 5 min three times and incubated with secondary antibody in Western Blocker and Signal Enhancer (MP) for 1 hour at room temperature. Then, membranes were washed with PBS + 2vol/vol% Tween 20 for 5 min three times and incubated with Immobilon ECL Ultra Western HRP Substrate (Millipore) for 5 min, then scanned in a Chemidoc MP imager (BioRad). For the primary antibodies, 1/1000 of anti-Rad21 from rabbit (Abcam: ab922), 1/1000 of anti-STAG1 from rabbit (proteintech: 14015-1-AP), 1/1000 of anti-STAG2 from rabbit (proteintech: 19837-1-AP), 1/1000 of anti-STAG2 from rabbit (proteintech: 19837-1-AP), 1/1000 of anti-M18BP1/KNL2 from rabbit (proteintech: 26264-1-AP), 1/10000 of anti-α-Tubulin from mouse (Sigma: 9026). For secondary antibody, HRP conjugated antibodies were used.

### Cell counting

To count the number of DLD-1 cells, the culture medium was removed using an aspirator. Then, 2.5 g/liter of Trypsin 1 mmol/liter EDTA solution (Gibco) was added and incubated for 5 min at 37°C. The same amount of complete culture medium as Trypsin was added to stop trypsinization. The cell solution was mixed with the same volume of 0.4 wt/vol% Trypan Blue Solution, and cell numbers were counted using Countess II.

### Cell fixation and immunofluorescence

Cells were fixed with 4% PFA (Electron Microscopy Services) in PBS for 10 min at room temperature. For salt pre-extraction, cells were incubated with 0.5 M NaCl for 30 min at room temperature, then the buffer was removed and the cells were washed with PBS and fixed with PFA. For detergent pre-extraction, cells were incubated with PBS + 0.5vol/vol% Triton X-100 for 60 sec at room temperature, then the buffer was removed, washed with PBS, and fixed with PFA.

Fixed cells were permeabilized by incubation in PBS + 0.5vol/vol% Triton X-100 for 15 min at room temperature, then cells were washed with PBS + 2vol/vol% Tween 20. The following antibodies were used for immunofluorescence: human anti-CREST (Immunovision HCT-100) and rabbit anti-RNA polymerase II CTD repeat YSPTSPS (phospho S2) (Abcam, ab5095). Cells were incubated with the primary antibody for 1 hour at 37°C. Cells were washed 3 × 5 min with PBS + 2vol/vol% Tween 20 and then incubated with secondary antibody for 1 hour. Cells were washed 3 × 5 min with PBS + 2vol/vol% Tween 20. Cells were postfixed with 4% PFA in PBS for 10 min at room temperature. 5EU-RNA was detected essentially as described (Sharp et al., 2020).

### Image acquisition and analysis

All images were acquired using a Nikon A1R confocal microscope equipped with a 60 × 1.4NA Vc lens and laser lines at 405, 488, 562, and 647 nm. Images were collected using a scan zoom of 2× or 4× (140 or 70 nm/pixel) using a galvano scanner. Images were acquired as Z stacks spaced 0.2 µm apart. Each experiment was repeated in three biological replicates. The fluorescence intensity of RNA Pol II PS2, CENP-A-SNAP-TMR, or GFP-AID-KNL2 signal was measured in the nucleus of each cell using a script in Fiji. For RNA Pol II analysis, one dot indicated one cell’s signal intensity in each nuclear area. For CENP-A or KNL2 analysis, one dot indicated one centromere’s signal intensity per centromere area. For each experiment, fluorescence intensity was normalized to the average intensity of untreated control cells. Data are shown as mean ± SD of three independent assays. A two-tailed unpaired t-test with the mean of the experiment was used to compare conditions.

## Acknowledgement

We thank the members of the Blower lab for their suggestions and contributions to this paper. We also thank Dr. Aaron Straight for providing the KNL2 tagging plasmid. This work was supported by funding from the NIH/National Institute of General Medical Sciences to M.D.B. (5RO1-GM122893), the Overseas Research Fellowships of the JSPS/Japan Society for the Promotion of Science to R.W., and the Overseas postdoctoral fellowships of the Uehara Memorial Foundation to R.W.

## Declaration of Interests

The authors declare no competing interests.

---

**Supplemental Figure 1.**
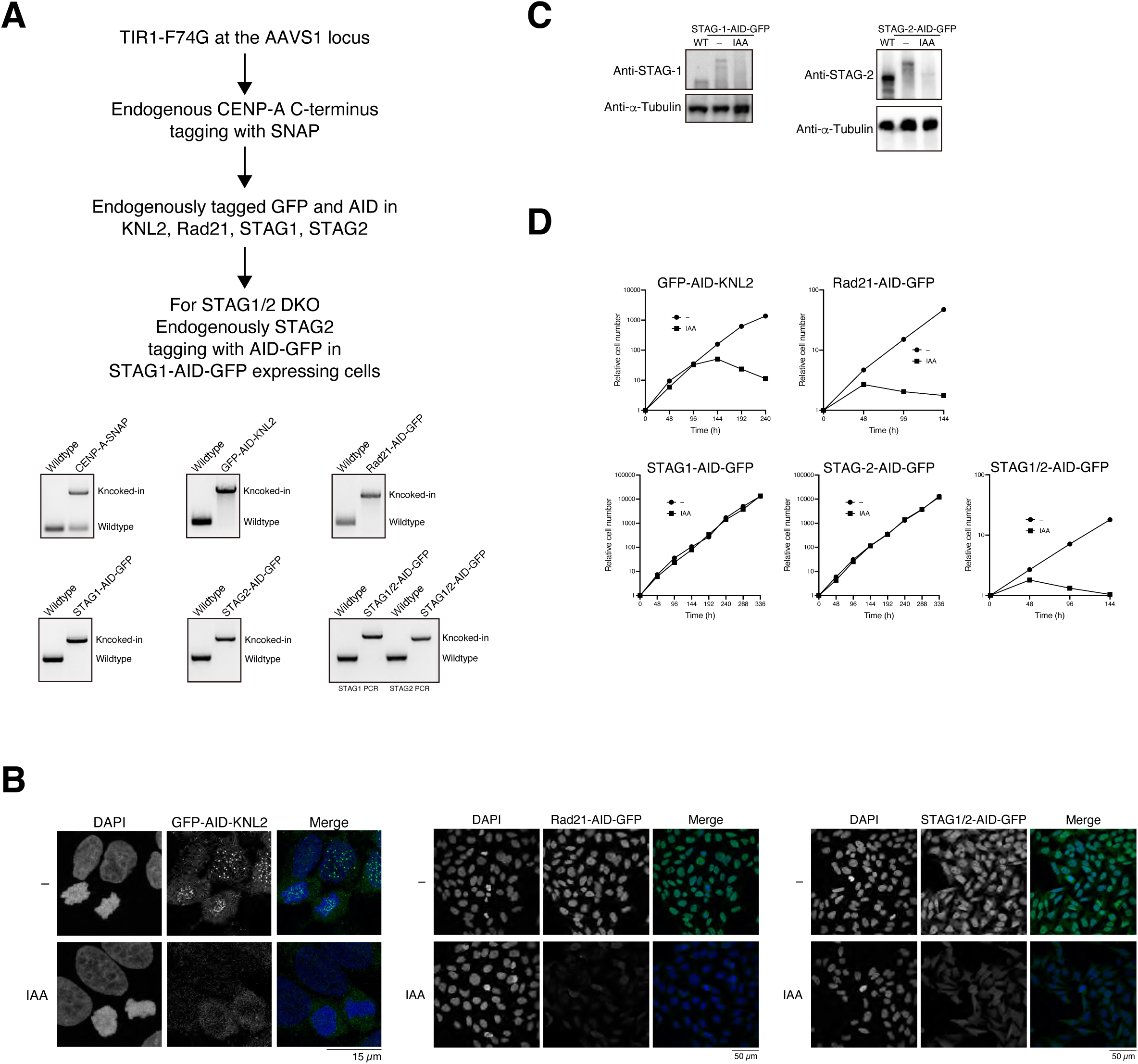
The auxin-inducible degron 2 (AID2) provided rapid depletion of targeting proteins in Human cells, together with growth defects. (A) Schematic representation of the CRISPR-Cas9-mediated knocking-in strategy in DLD1 cells. Images showed genomic PCR to confirm knocking-in to endogenous alleles. (B) Representative images of the protein signals endogenously tagged with AID and GFP. Those cells were cultured in the presence or absence of 5-Ph-IAA (IAA) for 1 hour and then fixed. DNA was stained with DAPI (blue). (C)The protein levels of endogenous STAG1 or STAG2, which were tagged with AID and GFP in human DLD1 cells constitutively expressing TIR1-F74G (right). AID-tagged protein was conditionally depleted by 5-Ph-IAA (IAA) addition for 2 hours. (D) Growth of DLD1 cells expressing AID-tagged proteins. Cell proliferation was examined in the presence or absence of IAA. Time shown represents hours after IAA addition.

**Supplemental Figure 2.**
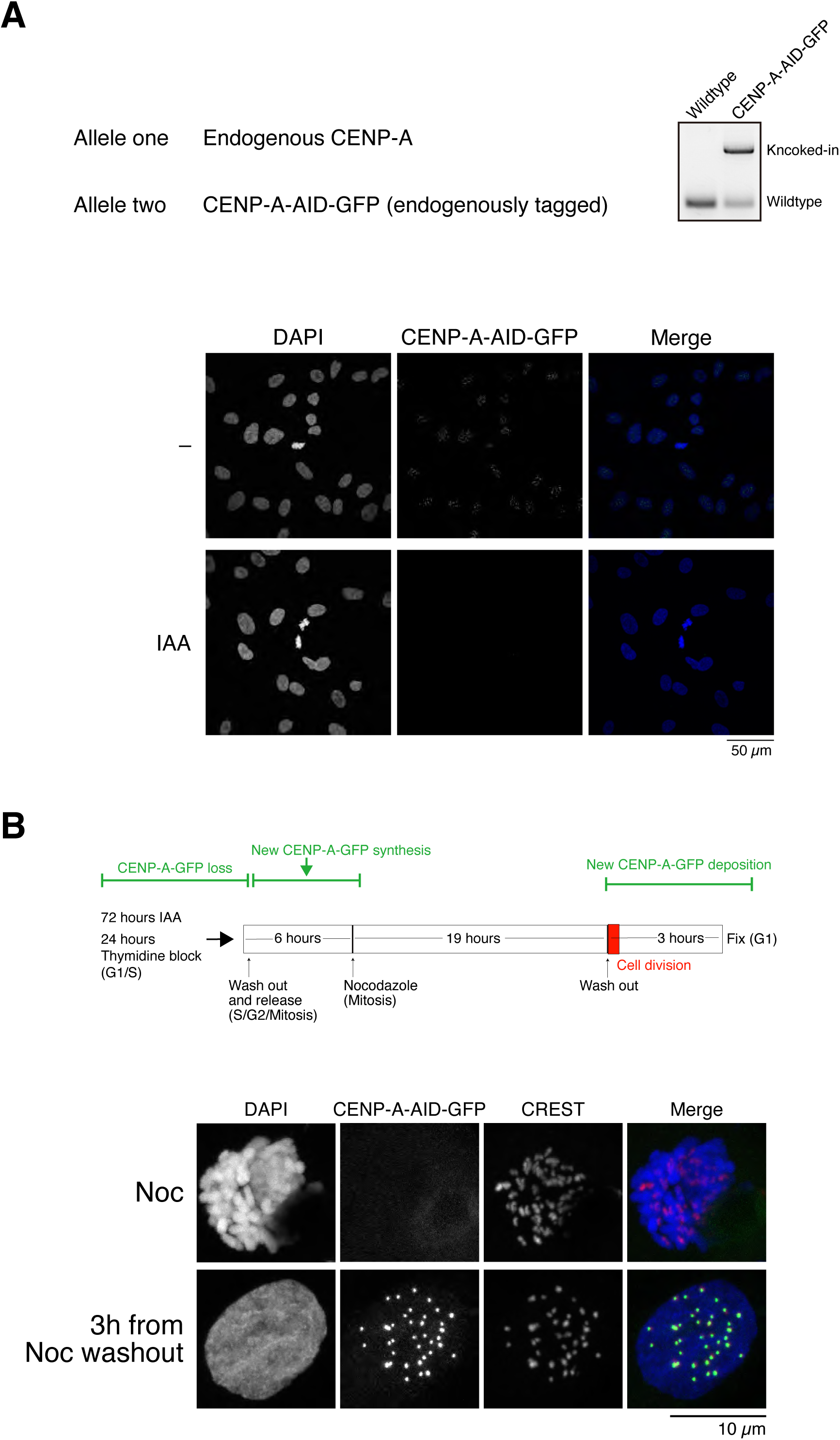
IAA washed out allowed CENP-A-AID-GFP recovery and deposition in newly G1 cells. (A) Image showed genomic PCR to confirm knocking-in to endogenous alleles in HeLa cells. Representative images of the CENP-A-AID-GFP signals from one allele. Those cells were cultured in the presence or absence of 5-Ph-IAA (IAA) for 1 hour and then fixed. DNA was stained with DAPI (blue). (B) Schematic representation of the experiment (top). Representative images of the CENP-A-AID-GFP signals. Those cells were cultured in the Nocodazol (Noc) overnight or washed out of it for 3 hours and then fixed. DNA was stained with DAPI (blue). The experiment followed the schematic representation of Figure 2A.

**Supplemental Figure 3.**
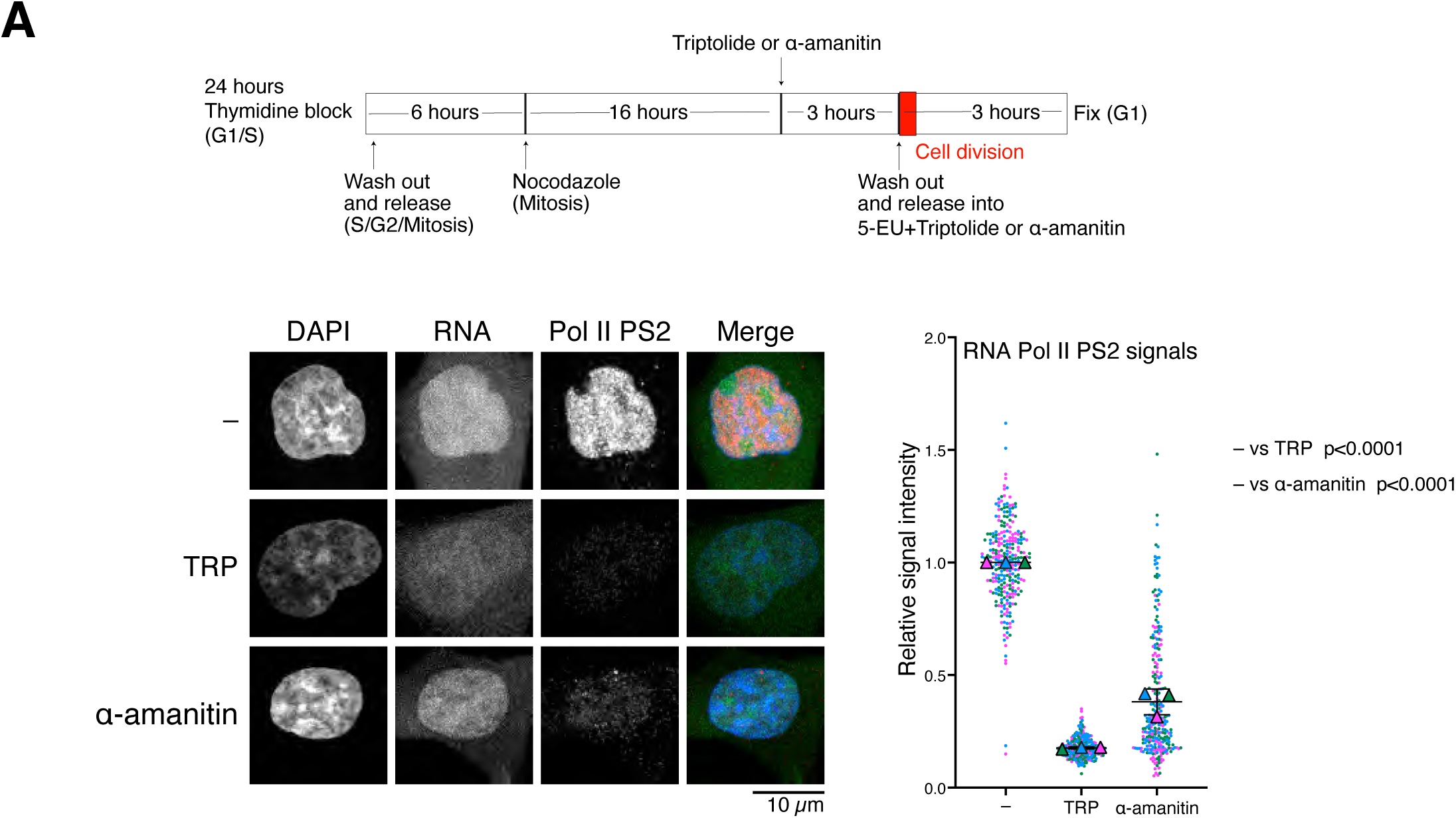
Triptolide and α-amanitin inhibited RNA pol II elongation in newly G1 cells. (A) Schematic representation of the experiment (top). Signals of RNA pol II CTD phospho-Ser2 (red) and EU-RNA (green) during G1 phase at 6 hours after Triptolide (TRP) or α-amanitin. DNA was stained with DAPI (blue).

**Supplemental Figure 4.**
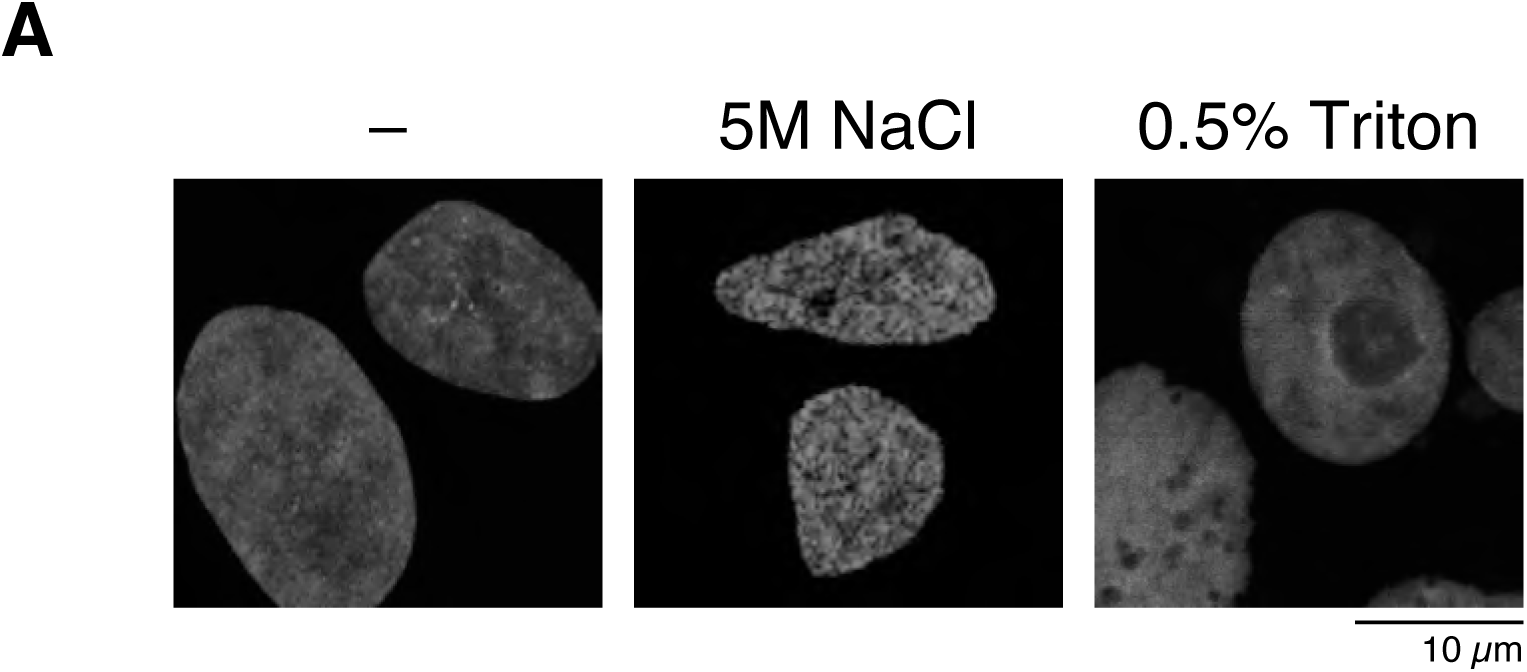
Salt extraction or detergent treatment resulted in chromatin conformation change. (A) DAPI image of HeLa cells treated with 0.5M NaCl for 30 min or 0.5% Triton-X for 60 sec before fixing, then the cells were fixed.

## Notes

### Competing Interest Statement

The authors have declared no competing interest.

